# Orthogonal tRNA Expression using Endogenous Machinery in Cell-Free Systems

**DOI:** 10.1101/2022.10.04.510903

**Authors:** Kosuke Seki, Joey L. Galindo, Michael C. Jewett

**Affiliations:** Department of Chemical and Biological Engineering, Northwestern University, Evanston, IL 60208, USA; Center for Synthetic Biology, Northwestern University, Evanston, IL 60208, USA; Chemistry of Life Processes Institute, Northwestern University, Evanston, IL 60208, USA

## Abstract

A wide variety of non-canonical amino acids (ncAAs) can be incorporated into proteins through the coordinated action of a stop codon suppressing tRNA and aminoacyl-tRNA synthetase. However, methods to discover and characterize suppressor tRNAs are generally lacking. In this work, we show that cell-free systems can express functional suppressor tRNAs using endogenous machinery and characterize their activity. This method is compatible with widely used orthogonal tRNAs, such as the *Methanocaldococus jannaschii* tyrosyl tRNA, the *Methanosarcina barkeri* pyrrolysyl tRNA, the *Methanomethylophilus Alvus* pyrrolysyl tRNA, and an engineered *Int* pyrroysyl tRNA. Modifying the workflow to evaluate TAA suppression revealed that the *M. jannaschii* and *M. alvus* are highly functional TAA suppressors in cell-free systems. Finally, we show that we can express two distinct tRNAs simultaneously, enabling the incorporation of multiple, distinct ncAAs. In total, our work shows that cell-free systems are useful platforms to express and characterize tRNAs.

## Introduction

Proteins are an extraordinary class of biomolecules that serve a broad range of functions due to the order and chemistry of the canonical twenty amino acids. While there have been many advances in protein engineering methods with the canonical amino acids to evolve new functions (1), proteins with non-canonical amino acids (ncAAs), those outside the natural twenty, remain cumbersome to synthesize and engineer (2, 3). Yet, these proteins have broad potential, as exemplified in recent works showing that ncAAs can modulate enzyme catalysis (4, 5), antibody binding (6, 7), material properties (8–10), and a variety of other important protein functions (2, 3).

A key challenge in ncAA incorporation is the discovery and characterization of tRNAs and aminoacyl-tRNA synthetases (aaRS – specific aaRSs are noted by their three letter amino acid code, i.e. alaRS for alanyl-tRNA synthetase) (11, 12). These pairs, called Orthogonal Translation Systems (OTS), must satisfy stringent requirements, including orthogonality towards endogenous tRNAs and aaRSs but not towards auxiliary translation factors like elongation factors and ribosomes, and sufficient aminoacylation activity to drive protein synthesis. Recent works have discovered several OTSs that satisfy these criteria, including the *Methanocaldococcus jannaschii* tyrRS:tRNA pair and several pyrrolysyl (pyl) tRNA:pylRS systems, which are mutually orthogonal (13–17). This has paved the way for exciting opportunities in the synthesis of proteins containing multiple, distinct ncAAs (18–21). However, laborious and time-consuming workflows in living cells impedes new discovery and characterization of OTS components.

Cell-free protein synthesis (CFPS) has potential to facilitate discovery and characterization of new components of OTSs. CFPS has already been used to efficiently synthesize proteins containing multiple instances of a ncAA (22, 23), incorporate a wide variety of ncAAs, including those that are toxic, impermeable across the cell membrane, or insoluble (24–26), incorporate multiple, distinct ncAAs (27), reassign sense codons (28–31), and screen a library of engineered aaRS variants (32). These advances were enabled by a combination of factors, including high volumetric productivity (> mg/mL protein), rapid protein synthesis (< 8 hours), scalability between volumes (nL to 100-L), the ability to bypass cellular constraints, and compatibility with high-throughput liquid handling systems (23, 33–35).

However, one challenge in cell-free systems is the expression of stop codon suppressing tRNAs. Expression of ncAA-containing proteins in CFPS is typically accomplished by using cell extracts that are enriched with suppressor tRNAs by overexpression *in vivo*, or by supplementing reactions with tRNAs purified by traditional *in vitro* transcription methods, which are both low-throughput methods. One work has shown that the *M. jannaschii* tyrosyl tRNA can be co-transcribed alongside a protein of interest using T7 RNA polymerase during the CFPS reaction (36). This system uses a linear DNA template called a ‘transzyme’ consisting of a buffer sequence, a T7 promoter, a hammerhead ribozyme for transcription of unfavorable 5’ sequences, and a tRNA sequence whose 3’ end terminates the DNA template. Besides this successful example of *in vitro* transcription of *M. jannaschii* tyrosyl tRNA, there is generally a lack of methods to evaluate TAG suppressors in CFPS.

In this work, we present a general cell-free method to enable expression and evaluation of stop codon suppressing tRNAs. We first designed and optimized endogenously driven constructs enabling tRNAs to be co-transcribed with mRNA templates for ncAA-containing proteins of interest. The tRNA construct contains a *proK E. coli* promoter and terminator on a plasmid template, allowing for endogenously driven tRNA expression and maturation. We show that this method is generalizable for expression and evaluation of a panel of commonly used TAG- and TAA-suppressing tRNAs that were chosen for their mutual orthogonality (13, 14, 37). Finally, we use them to incorporate multiple, distinct ncAAs into a single protein by *in situ* transcription of two separate tRNAs. We anticipate that this cell-free workflow will enable future discovery and characterization of tRNAs that may be candidates for novel OTSs.

## Methods

### Reagents and Materials

All materials were purchased from Sigma-Aldrich unless otherwise stated. Q5 High-Fidelity DNA Polymerase, DpnI, DNA Loading Dye, T5 Exonuclease, Phusion Polymerase, ET SSB, Taq DNA Ligase, and GamS were purchased from New England Biolabs. Agarose was purchased from Invitrogen. SYBR Safe Dye was purchased from APExBio. DNA was synthesized by either IDT or Twist Biosciences and is specified when described. Nuclease-Free Water was from Ambion. DH10β chemically competent cells, BL21 (DE3) Star chemically competent cells, Slide-a-Lyzer Dialysis Casettes, Pierce Protein Concentrators, NuPAGE LDS Sample Buffer, 4-12% Bis-Tris NuPAGE gels, and MES Buffer were purchased from ThermoFisher. AcquaStain Protein Gel was purchased from Bulldog Bio. Overnight Express TB Medium, Benzonase, and Amicon Ultra 0.5 mL Centrifugal Filters were purchased from Millipore. Ammonium Glutamate was purchased from MP Bio. Total *E. coli* tRNA and Phosphoenolpyruvate were purchased from Roche. Bpy and AzK were purchased from Toronto Research Chemicals. ^14^C leucine, Filtermat A, and Meltilex A were purchased from Perkin Elmer. Streptactcin XT Spin Columns were purchased from IBA Life Sciences.

### DNA Amplification

All PCRs were conducted using Q5 High-Fidelity DNA Polymerase following the manufacturer’s protocols. Primers and gBlocks were synthesized by IDT. Products from all PCRs were analyzed by agarose gel electrophoresis. Agarose gels were prepared by dissolving agarose in TAE buffer at 1% w/v (for tRNA plasmid backbone amplification and PJL1 backbone amplification) or at 2% w/v (for transzyme amplification) and stained with SYBR Safe dye. 2.5 μL of PCR product was mixed with 2.5 μL of water and 1 uL of 6X Loading Dye and loaded into the appropriate agarose gel. 1% w/v agarose gels were run at 120 V for 30 minutes, and 2% w/v agarose gels were run at 120 V for 1 hr. Gels were imaged using a Bio-Rad Gel Doc XR+ to confirm amplicon size.

### DNA Assembly

tRNA expression plasmids were assembled using Gibson Assembly. tRNA expression plasmid backbone was amplified as previously described under DNA Amplification. PCR reactions were column purified using Zymo DNA Clean and Concentrate and resuspended in 10 μL of Nuclease-Free water. DNA was digested by adding 5 μL of 10 X CutSmart Buffer, 34 μL of Nuclease-Free water, and 1 μL of DpnI to the purified PCR product. The mixture was incubated at 37 °C for one hour followed by 80 °C for 20 minutes, and then column purified again. The Gibson assembly reaction was prepared by mixing the tRNA plasmid backbone and an appropriate gBlock containing the tRNA in a 1:3 molar ratio, respectively, along with a homemade 3 X Gibson Assembly Master Mix (75 mM Tris-HCl pH 7.5, 7.5 mM MgCl_2_, 0.15 mM dNTPs, 7.5 mM DTT, 0.75 mM NAD, 0.004 U/μL T5 Exonuclease, 0.025 U/μL Phusion Polymerase, 4 U/μL Taq DNA Ligase, and 3.125 μg/mL ET SSB) and an appropriate volume of water. This mixture was incubated at 50 °C for one hour. 3 μL of the Gibson Assembly reaction was transformed into chemically competent DH10β cells following manufacturer’s instructions. Cells were streaked onto LB-agar plates containing 50 μg/mL kanamycin and incubated overnight at 37 °C. Single colonies were inoculated into 5 mL of LB-Miller media with 50 μg/mL kanamycin and plasmid DNA was isolated using ZymoPURE Miniprep kits. Accurate assembly of the plasmid was confirmed by Sanger Sequencing at GeneWiz using primers PJL1-Forward (ctgagatacctacagcgtgagc) and PJL1-Reverse (cgtcactcatggtgatttctcacttg).

The 1TAA-sfGFP and 1TAG-1TAA sfGFP were similarly assembled using Gibson Assembly. The PJL1 backbone was amplified by PCR. The insert and backbone were assembled, transformed, and inserts were sequence verified as described above. Sanger Sequencing was done by GeneWiz using their T7-forward and T7-reverse primers.

### DNA Purification

Plasmids for all CFPS reactions were purified by ZymoPURE Midiprep kits. 50 mL cultures were prepared using LB-Miller media containing kanamycin at 50 μg/mL and were inoculated from single colonies on a plate or from a glycerol stock. DNA was purified following the manufacturer’s instructions and was verified by Sanger Sequencing as described above.

For transzyme constructs, PCR products were analyzed by agarose gel electrophoresis and column purified using Zymo DNA Clean and Concentrate to ∼ 300 ng/μL.

### Cell Extract Preparation

759.T7 is a genomically engineered *E. coli* strain used in all cell-free experiments in this work. This strain has several features, including (1) all TAG codons are recoded to TAA, (2) Release Factor 1 was knocked out, (3) Negative effectors of CFPS were knocked out, and (4) T7 RNA Polymerase was integrated onto its genome (22, 23).

A 150 mL overnight culture was inoculated from a glycerol stock and incubated at 34 °C overnight with 250 RPM shaking. The next day, the OD_600_ of a 10 X dilution of the overnight culture was measured using a NanoDrop 2000C in a 1 mL cuvette. This was used to inoculate 10 L of 2xYPTG (16 g/L tryptone, 10 g/L yeast extract, 5 g/L NaCl, 7 g/L KH_2_PO_4_, and 3 g/L K_2_HPO_4_) at OD_600_ = 0.075 in a 10 L Sartorius Biostat Cplus fermenter at 34 °C. At OD_600_ = 0.5, 10 mL of 1 M IPTG (final concentration = 1 mM) was added to the culture to induce expression of T7 RNA polymerase. Cells were grown until OD_600_ = 3.0. Cells were then harvested by centrifugation (Beckman-Counter Avanti J-26) at 5,000 x g for 15 minutes at 4 °C, washed by resuspending with S-30 Buffer (10 mM Tris-Acetate pH 8.2, 14 mM Mg Acetate, 60 mM K Acetate, 2 mM DTT) and centrifugation for a total of three washes. Cells were pelleted by a final centrifugation at 10,000 x g for 5 min at 4 °C. Cells were then flash frozen at -80 °C until lysis.

Cells were thawed on ice and resuspended in 0.8 mL S-30 buffer / g wet cell mass by vortexing. Cells were lysed by sonication using a Q125 Sonicator (QSonica) at 50 % amplitude using 3 × 45s on and 59 s off cycles to a total of 950 J. Lysate was centrifuged at 12,000 x g for 10 minutes at 4 °C to remove insoluble debris. The supernatant was collected and incubated at 37 °C with 250 RPM shaking in a run-off reaction. The lysate was re-centrifuged at 10,000 x g for 10 min at 4 °C to remove insoluble components that appeared after run-off. Finally, the clarified lysate was dialyzed against 200 volumes of S-30 buffer in a 10 kDa MWCO Slide-a-Lyzer Dialysis Casette. Lysate was then aliquoted into single-use aliquots, flash frozen in liquid nitrogen, and stored at -80 °C.

### aaRS Purification

pET.BCS-BpyRS was cloned as previously described using Gibson Assembly (38). Expression plasmids for the Chimeric pylRS, the *M. alvus* pylRS, and the *Lum1* pylRS were synthesized in the pET.BCS backbone by Twist Biosciences. All aaRSs contained a C-terminal His-tag.

Plasmids were transformed into chemicomp BL21 (DE3) Star following the manufacturer’s instructions. Cells were plated onto LB-agar plates containing 100 μg/mL carbenicillin and incubated overnight at 37 °C. A single colony was picked into 5 mL of LB-Carb and grown overnight at 37 °C with 250 RPM shaking. 0.25 mL of expression culture was inoculated into 250 mL of Overnight Express TB Medium containing carbenicillin and incubated for ∼16 hours at 37 °C with 250 RPM shaking. The next day, expression cultures were pelleted by centrifugation at 5,000 x g for 10 min at 4 °C, washed with Buffer 1 (300 mM NaCl, 50 mM Sodium Phosphate Monobasic pH 8) supplemented with 10 mM imidazole pH 8, pelleted again, and then stored at -80 °C.

Cells were thawed on ice and resuspended in 3 mL Buffer 1 supplemented with 10 mM imidazole pH 8 and benzonase / g wet cell mass. The cell solution was passed through a syringe needle and lysed by homogenization (Avestin B3) at ∼20,000 PSI. Insoluble components were then pelleted by centrifugation at 20,000 x g for 15 minutes at 4 °C. Supernatant was removed and incubated with 5 mL of pre-equilibrated Ni-NTA resin (Qiagen) for one hour at 4 °C with gentle shaking. The resulting suspension was centrifuged at 3,800 x g for 5 minutes at 4 °C, and supernatant was carefully decanted. The resin was washed five times with 20 mL of Buffer 1 + 20 mM imidazole pH 8 in a 50 mL falcon tube in batches. Finally, the resin was packed into a column and proteins were eluted from the column using 20 mL of Buffer 1 + 0.5M imidazole pH 8. Elution fractions containing protein, as measured by A280 on Nanodrop 2000c, were combined and dialyzed overnight into 2 L of Buffer 1 + 40 % v/v glycerol with gentle stirring at 4°C using a Slide-a-Lyzer Dialysis Casette (3.5 kDa MWCO). The dialysis buffer was changed after an overnight dialysis and dialyzed for an additional 4 hours at 4 °C. Samples were then removed from the dialysis cassette, concentrated using a Pierce Protein Concentrator (3K MWCO), measured by Nanodrop using extinction coefficients and molecular weights calculated by ExPasy ProtParam, flash frozen in single use aliquots, and stored at - 80 °C.

Protein purity was analyzed by SDS-PAGE. 1 μL of concentrated protein was mixed with 3.75 μL of NuPAGE 4X LDS Sample Buffer, 1.5 μL of 1 M DTT, and water to 15 μL. This sample was denatured at 95 °C for 10 minutes. Samples were loaded onto a 4-12 % Bis-Tris NuPAGE gel and run in 1X MES Buffer for 45 minutes at 180 V. Gels were stained with gentle shaking using AcquaStain Protein Gel for 15 minutes, destained in water for 1 hour, and then imaged.

### CFPS

CFPS reactions are based on a modified PANOx-SP system. Generally, 5 μL CFPS reactions were set up in 0.2 mL PCR tubes in conditions of 4-12 mM Magnesium Glutamate, 10 mM Ammonium Glutamate (MP Bio), 130 mM Potassium Glutamate, 1.2 mM ATP, 0.85 mM of GTP, CTP, and UTP each, 0.03 mg/mL Folinic Acid, 0.17 mg/mL tRNA (Roche), 0.4 mM NAD, 0.27 mM CoA, 4 mM Oxalic Acid, 1 mM Putrescine, 1.5 mM Spermidine, 57 mM HEPES pH 7.2, 33 mM Phosphoenolpyruvate (Roche), and 30 % v/v 759.T7 cell extract. The optimal Magnesium Glutamate concentration was determined for each batch of cell extract. PJL1-sfGFP and the tRNA expression plasmids were added at 5 ng/μL and 20 ng/μL, respectively, though data in **Figure 2b** and **2d** were generated using the traditional 13.33 ng/μL for both plasmids. Bpy and AzK (Toronto Research Chemicals) were added at final concentrations of 1 mM, along with 1 mg/mL of the appropriate aaRS (**Table 1**). For linear templates, transzyme constructs were added at a final concentration of 30 ng/μL, WT sfGFP linear template was added in an equal volume as PJL1-sfGFP plasmid template, and GamS was added at 1 μM. Reactions were incubated either at 30 °C or 37 °C in an incubator, or, for **Figure 2d**, were conducted in a thermocycler with a temperature gradient.

**Table 1:** Description of OTSs and ncAAs used in this study

| tRNA | ncAA | aaRS |
| --- | --- | --- |
| *M. jannaschii* tyr tRNA | Bipyridyl alanine (Bpy) | BpyRS (44) |
| *M. barkeri* pyrrolysyl tRNA | Azidolysine (AzK) | IPYE Chimeric PylRS (45) |
| *M. alvus* pyl tRNA (13) | AzK | *M. alvus* PylRS (13) |
| <sup>A17</sup> <sup>V</sup>C10 *Int* pyl tRNA (14) | AzK | *Lum1* (14) |

### CFPS Quantification of sfGFP

sfGFP concentration in CFPS was quantified by using a standard curve relating sfGFP fluorescence to protein concentration as measured by ^14^C scintillation counting. WT sfGFP with ^14^C leucine was synthesized by adding ^14^C leucine at a final concentration of 10 μM in 15 μL CFPS reactions. CFPS reactions were spun down at 21,000 x g for 15 minutes at 4°C, and 5 μL of supernatant was removed to isolate soluble proteins. An equal volume of 0.5 M KOH was added to the soluble proteins and incubated at 37 °C for 20 minutes. 4 μL of each sample were than spotted onto two Filtermat A fiberglass paper sheets and allowed to dry under a heat lamp. One of the filtermats was washed three times with 5% w/v TCA for 15 min at 4 °C, and then washed with 100% ethanol at room temperature for 15 min. The filtermat was again allowed to dry under a heat lamp. Meltilex A was then applied to both filtermats. After cooling and solidifying, scintillation counts were measured using the MicroBeta^2^ (Perkin Elmer) and soluble yields were calculated as previously described (39).

Dilutions of the sfGFP were made in 1X PBS and measured on a plate reader (BioTek Synergy 2). A standard curve was built using a linear regression to fit fluorescence measurements to protein yields as calculated by ^14^C scintillation counting.

### Preparation of Proteins for intact protein ESI-MS

Proteins were synthesized using CFPS as described previously, except that reactions were scaled up to 75 or 150 μL CFPS reactions in a 50 mL falcon tube. Proteins were purified from CFPS reactions using Strep-Tactin XT Spin Columns according to the manufacturer’s instructions. Proteins were then buffer exchanged into 0.1 M Ammonium Acetate using Amicon Ultra 0.5 mL Centrifugal Filters (10 kDa MWCO).

Proteins were analyzed with ESI-MS as described in previous publications with slight modifications (40). 8 μL of 2 μM purified, buffer-exchanged protein (16 pmol) were injected into a Bruker Elite UPLC coupled to a Bruker Impact II UHR TOF Mass Spectrometer. Proteins were deconvoluted using a m/z range of 20,000 to 30,000 Da.

## Results

In developing a general method for tRNA expression *in vitro*, we first evaluated whether transzymes could transcribe orthogonal tRNAs during CFPS (**Figure 1a**). Linear DNA templates for the *M. jannaschii* tyrosyl tRNA, the *M. barkeri* pyl tRNA, the *M. Alvus* pyl tRNA, and an engineered ^A^17 ^V^C10 *Int* pyl tRNA were PCR amplified, purified, and added into CFPS reactions at 30 ng/μL, a concentration previously shown to be optimal (36). These CFPS reactions consist of 759.T7 extract made from an *E. coli* strain lacking TAG codons and Release Factor 1 (RF1) (23), GamS to enable expression of linear DNA templates (**Figure 1b**) (41), an appropriate ncAA, aaRS, and tRNA (listed in **Table 1**), traditional CFPS reagents (42), and a 1TAG-sfGFP reporter containing a premature amber codon at T216. We found that all transzymes supported 1TAG-sfGFP expression (**Figure 1c**). Interestingly, 1TAG-sfGFP expression using the *M. jannaschii* tRNA template did not require GamS for expression, and yields using the *M. alvus* tRNA were decreased. While successful, this method generally faces several drawbacks, including the necessity to design unique hammerhead ribozyme sequences for each tRNA (43), the need for GamS, and the potential for inefficient cleavage of tRNAs leading to low protein yields.

Alternatively, we hypothesized that using endogenous *E. coli* machinery for tRNA transcription and maturation could be a better approach because it bypasses the need for hammerhead ribozymes (**Figure 2a**). The *proK* promoter and terminator sequences are often used for orthogonal tRNA expression *in vivo* (46, 47), but it was unclear if these sequences would be functional for tRNA transcription and processing in cell extracts. To test this, we first cloned the *M. jannaschii* tyr tRNA flanked by the *proK* promoter and terminator on a plasmid. CFPS reactions containing 13.33 ng/μL of both the tRNA and 1TAG-sfGFP plasmid were set up and incubated at 30 °C. We measured high yields of ∼725 μg/mL of 1TAG-sfGFP in the presence of both the tRNA and 1TAG-sfGFP plasmids, showing that tRNAs expressed with endogenous machinery are as efficient as transzyme constructs for amber suppression (**Figure 2b**). We also noticed that the addition of the 1TAG-sfGFP plasmid alone resulted in significant readthrough, which we define as non-specific suppression of the TAG codon (**Figures 2b, 1c**). Electrospray-Ionization Mass Spectrometry (ESI-MS) analysis of the intact protein suggests that readthrough occurs primarily due to non-specific incorporation of glutamine and tyrosine, both of which have been previously observed to suppress the TAG codon (**Figure 2c**) (48). Despite the readthrough, these results suggest effective amber suppression from transcription and maturation of tRNAs using endogenous machinery in CFPS.

**Figure 1:**
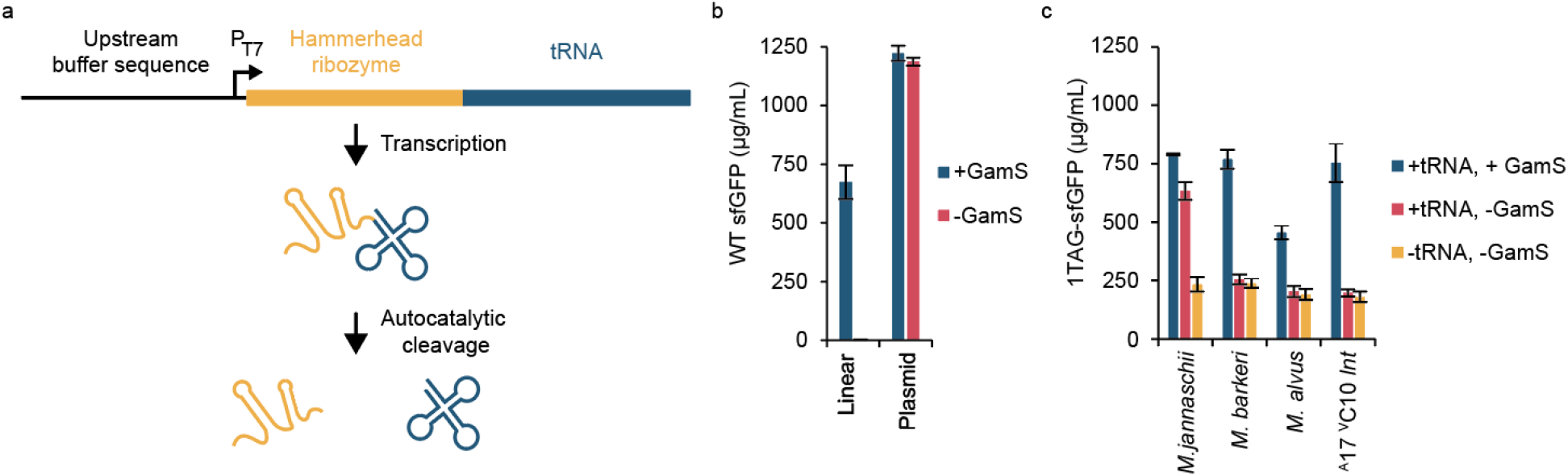
Linear transzyme constructs can express a panel of orthogonal tRNAs in CFPS. (**a**) Schematic of transzymes. A linear DNA template which encodes an upstream buffer sequence, a T7 promoter, a hammerhead ribozyme specific for a tRNA, and the orthogonal tRNA is added into CFPS. T7 RNA polymerase in CFPS transcribes the transzyme construct and the hammerhead ribozyme autocatalytically cleaves itself off to generate a mature tRNA. Expression of WT-sfGFP from a linear template requires GamS, while plasmid templates do not. (**c**) A panel of orthogonal tRNAs can be expressed in CFPS by supplementing the reaction with GamS. Reactions were done at 30°C.

**Figure 2:**
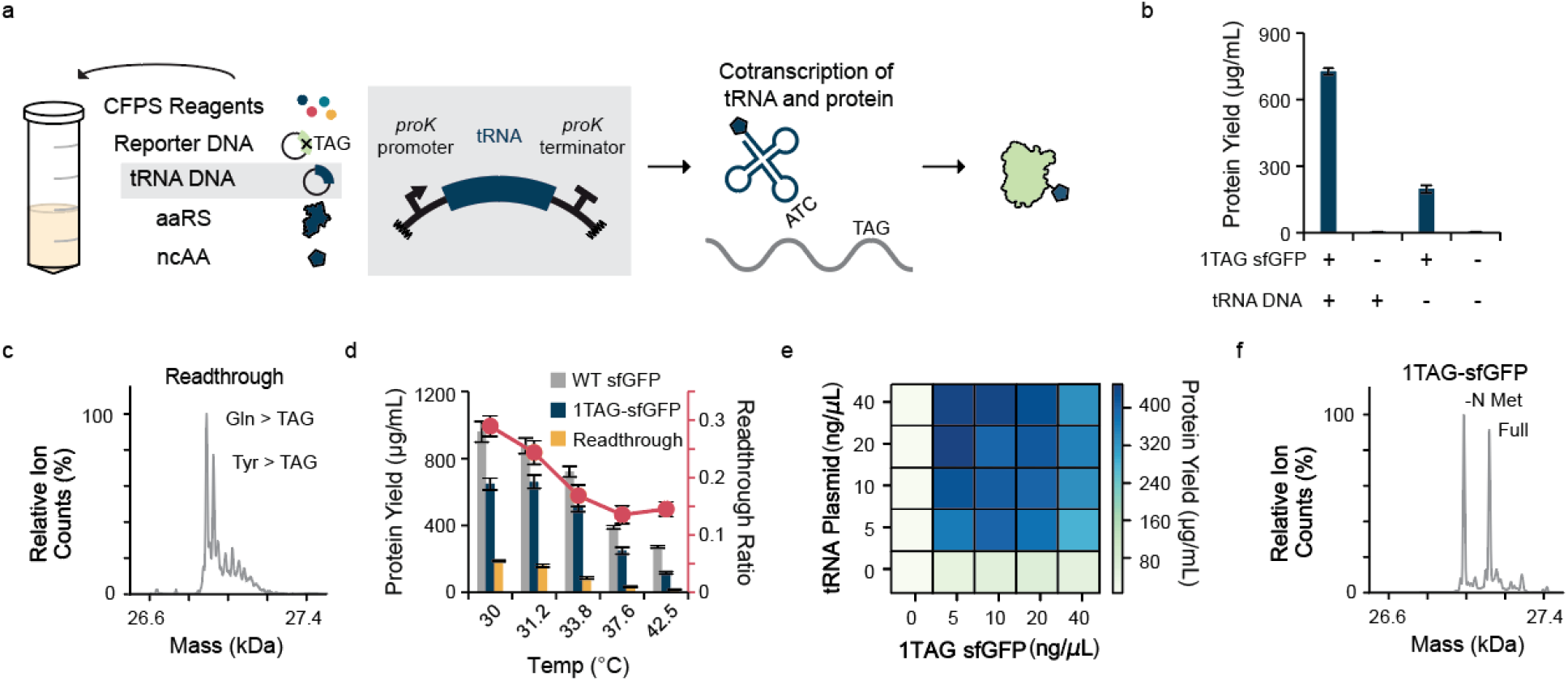
Expression of orthogonal tRNAs from *proK* based constructs is functional and can be optimized for efficient and accurate ncAA incorporation. (**a**) Schematic of orthogonal tRNA expression in CFPS. Orthogonal tRNAs are expressed from an endogenous *proK* promoter, which drives expression of a 1TAG-sfGFP reporter. (**b**) Expression of 1TAG-sfGFP is most efficient in the presence of both the 1TAG-sfGFP and tRNA plasmids. 1TAG-sfGFP expression in the absence of tRNA suggests readthrough by endogenous translation components. (**c**) ESI-MS analysis of readthrough products of 1TAG-sfGFP in the absence of tRNA reveals nonspecific incorporation of glutamine and tyrosine. Predicted masses: 26864.23 Da (WT sfGFP), 26892.24 Da (1TAG-sfGFP with glutamine readthrough), and 26926.30 Da (1TAG-sfGFP with tyrosine readthrough). Observed masses: 26890.37 Da, 26924.36 Da. All masses here include the excision of the N-terminal methionine, which is common in E. coli. (**d**) A titration of reaction temperature reveals that readthrough is inversely proportional to temperature. (**e**) Optimization of plasmid concentrations to reduce unfavorable competition between transcriptional and translational resources improve 1TAG-sfGFP synthesis. (f) ESI-MS analysis shows accurate ncAA incorporation (Predicted: 26988.22 (-N Methionine) and 27119.42 (Full), Observed: 26988.62 and 27118.65 Da).

We next aimed to reduce readthrough and optimize expression of 1TAG-sfGFP in this system. Hypothesizing that readthrough results from non-cognate base-pairing between the codon and anticodon in the absence of RF1, we increased the reaction temperature to discourage non-cognate base pairing. We found that readthrough and temperature are inversely proportional, and that the readthrough ratio, defined as the quotient between 1TAG-sfGFP yields in the absence and presence of tRNA DNA, was minimized at higher temperatures (**Figure 2d**). A reaction temperature of ∼37 °C, compared to the traditionally used 30 °C, balanced protein yields while reducing readthrough and was used for the remainder of TAG suppressing experiments (**Figure 2d**). To compensate for the loss in yield at 37 °C, we then optimized the ratio of the 1TAG-sfGFP and tRNA plasmids to reduce unfavorable competition (**Figure 2e**). A high concentration of tRNA template (∼20 ng/µL) and a low concentration of 1TAG-sfGFP template (∼5 ng/µL) produced the highest protein yields. The synthesized 1TAG-sfGFP product was analyzed by ESI-MS, which displayed a mass shift consistent with ncAA incorporation (**Figure 2f**). This shows that CFPS reactions can be optimized for relatively high-yielding ncAA incorporation within proteins.

After optimizing our system for tRNA co-expression and ncAA incorporation, we evaluated a panel of other commonly used orthogonal TAG- and TAA-suppressing tRNAs. We cloned a panel of orthogonal pyl tRNAs (**Table 1**) into the *proK* cassette and supplemented those plasmids into CFPS reactions. We observed consistent yields of 1TAG-sfGFP across each of the TAG-suppressing tRNAs in this expression cassette (**Figure 3a**). ESI-MS analysis shows efficient incorporation of AzK, suggesting tRNA expression and amber suppression (**Figure 3b**). We then tested the equivalent TAA-suppressing tRNAs using a 1TAA-sfGFP reporter, with TAA at the T216 residue. These reactions were run at 30 °C as we expected RF2 to prevent readthrough of TAA. Interestingly, TAA-suppressing tRNAs exhibited a range of activities (**Figure 3c**). The 1TAA-sfGFP yields using the *M. jannaschii* and *M. alvus* tRNAs were relatively high at ∼75-80% of their respective 1TAG-sfGFP yields. However, yields of 1TAA-sfGFP were 25% of the 1TAG-sfGFP yields using the *M. barkeri* tRNA, and were negligible for the ^A^17 ^V^C10 *Int* tRNA. ncAA incorporation at TAA by the *M. jannaschii, M. barkeri*, and *M. alvus* tRNAs was confirmed by ESI-MS analysis of purified proteins (**Figure 3d**). These data show that this method is generalizable towards other TAG-suppressing tRNAs and can identify functional TAA-suppressing tRNAs.

**Figure 3:**
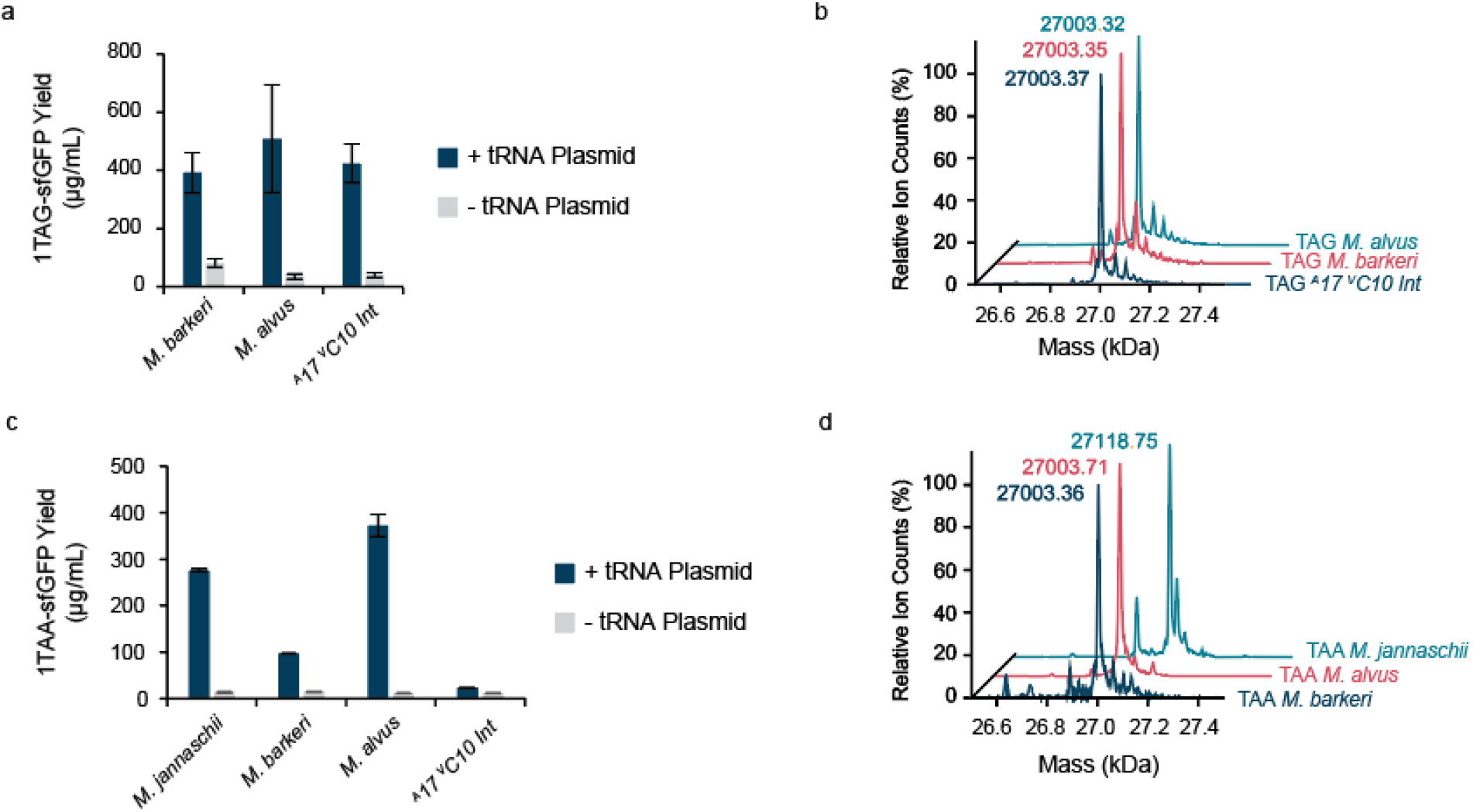
A panel of TAG and TAA suppressing orthogonal tRNAs can be expressed *in situ* during CFPS. (**a**) Commonly used pyrrolysyl tRNAs support incorporation of AzK. (**b**) The predicted mass for 1TAG-sfGFP with AzK at T216, 27004.25 Da, is consistent with the observed masses (masses include excision of the N-terminal methionine). Orthogonal tRNAs from *M*.*jannaschii, M. alvus*, and *M. barkeri* reprogrammed as TAA suppressors enable suppression of the 216TAA codon, with a range of observed activities. (**d**). TAA suppression is confirmed with intact-protein MS. The predicted mass for Bpy incorporation at T216, 27119.42 (including the N-terminal methionine), is consistent with the observed mass, and observed masses for AzK are consistent with the theoretical mass.

Finally, with several TAG- and TAA-suppressing tRNAs at hand, we aimed to demonstrate multiple, distinct ncAA incorporation within a single protein using two distinct codons. The *M. jannaschii* tyr-tRNA and the *M. alvus* pyl-tRNA were chosen to decode TAG and TAA, respectively, due to their mutual orthogonality, the anticodon-independent nature of pylRSs (49, 50), and the ability of *in situ* expressed *M. alvus* tRNAs to suppress TAA (**Figure 3c**). A 1TAG-1TAA-sfGFP reporter, containing TAG at T216 and TAA at N211, was successfully synthesized only when both tRNA templates were present (**Figure 4a**). Small amounts of readthrough were observed in the presence of only a single tRNA, with the TAA-suppressing tRNA generating slightly higher yields of readthrough product. This is consistent with literature suggesting that TAA-suppressing tRNAs can also readthrough TAG codons (19, 27, 51). ESI-MS of the product in the presence of both tRNAs displays mass shifts consistent with incorporation of each ncAA (**Figure 4b**). Thus, co-transcription of two mutually orthogonal tRNAs can direct the incorporation of two unique ncAAs within a single protein.

**Figure 4:**
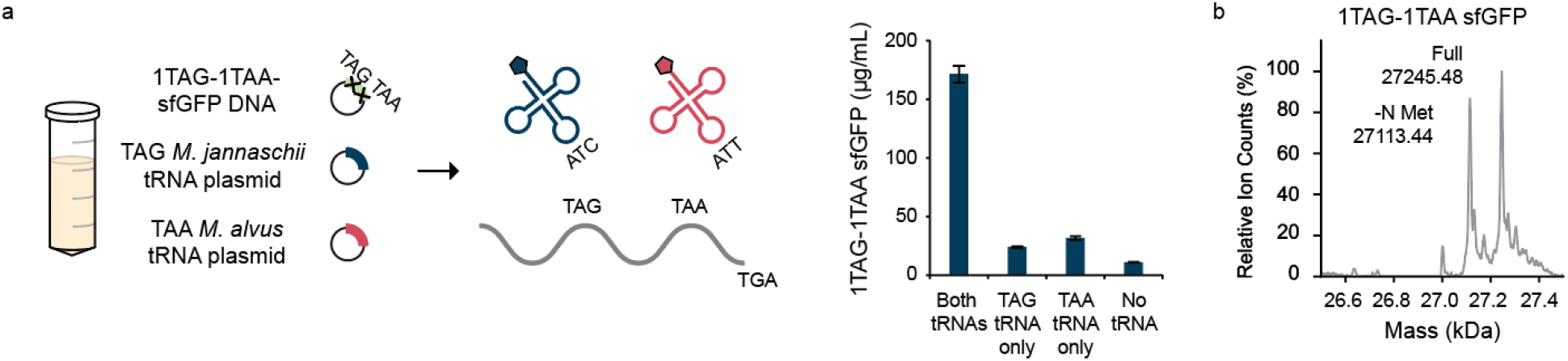
Incorporation of two, distinct ncAAs within a single protein. (**a**) Two mutually orthogonal tRNAs can be suppressed within a single protein. Both tRNAs are required for efficient full-length protein synthesis. (**b**) ESI-MS of 1TAG-1TAA sfGFP confirms incorporation of two, unique ncAAs (predicted: 27246.44 (full) and 27115.25 (-NMet)).

## Discussion

In this work, we developed a cell-free platform to express and evaluate stop codon suppressing tRNAs. This method co-transcribes a tRNA using endogenous *E. coli* machinery alongside an mRNA for a protein of interest to incorporate ncAAs. Our work expands on previous state-of-the-art technologies by showing that a wide variety of orthogonal tRNAs can be expressed *in situ* during CFPS, including TAG- and TAA-suppressing tRNAs, as well as both simultaneously.

We are exploring other lines of study to improve this system. Interestingly, we found that mutation of the anticodon from CTA to TTA decreased protein yields using the *M. barkeri* and ^A^17 ^V^C10 *Int* tRNAs. Certain factors are unlikely to explain this, such as reduced tRNA recognition by its cognate PylRS, as this process is independent of the anticodon (50), or RF2-based truncation at TAA, as other tRNAs were able to support reasonable yields of 1TAA-sfGFP (**Figure 3c**). We believe that factors related to the tRNA itself, such as expression, maturation, or folding, are likely responsible, though how the C > T mutation in the anticodon affects these is a topic of interest. Better understanding of the tRNA transcription and maturation process within this context will enable these reactions to be generalizable across a variety of tRNAs and codons, which is becoming increasingly important as sense codons become available for recoding (52, 53).

We anticipate that this technology to quickly express and characterize tRNAs will enable several exciting applications. Cell-free systems have unique advantages for tRNA expression and evaluation, including rapid protein synthesis and modularity. In addition, we imagine that the ability to scale CFPS reactions will enable high-throughput testing of other stop-codon suppressing tRNA sequences. In total, this cell-free tRNA expression platform enables simplified expression and characterization of stop codon suppressing tRNAs.

## Acknowledgements

The authors would like to acknowledge Dr. Andrew Hunt, Dr. Steve Fleming, Camila Kofman, Dr. Jessica Willi, Dr. Antje Krüger, and Dr. Jaeyoung K. Jung for helpful scientific discussions, Dr. Jasmine M. Hershewe for plasmid design, and Dr. Ashty Karim for help in manuscript preparation. This work was supported by the Army Research Office (W911NF-18-1-0200). This work made use of the IMSERC MS facility at Northwestern University, which has received support from the Soft and Hybrid Nanotechnology Experimental (SHyNE) Resource (NSF ECCS-2025633), and Northwestern University. We would like to thank Dr. Benjamin Owen and Dr. Fernando Tobias for training and assistance with mass spectrometry.

## Notes

### Competing Interest Statement

The authors have declared no competing interest.

## References

1. Wang, Y., Xue, P., Cao, M., Yu, T., Lane, S.T. and Zhao, H. (2021) Directed Evolution: Methodologies and Applications. Chem. Rev., 121, 12384–12444.

2. Zhang, W.H., Otting, G. and Jackson, C.J. (2013) Protein engineering with unnatural amino acids. Curr. Opin. Struct. Biol., 23, 581–587.

3. Dumas, A., Lercher, L., Spicer, C.D. and Davis, B.G. (2015) Designing logical codon reassignment - Expanding the chemistry in biology. Chem. Sci., 6, 50–69.

4. Burke, A.J., Lovelock, S.L., Frese, A., Crawshaw, R., Ortmayer, M., Dunstan, M., Levy, C. and Green, A.P. (2019) Design and evolution of an enzyme with a non-canonical organocatalytic mechanism. Nature, 570, 219–223.

5. Pott, M., Tinzl, M., Hayashi, T., Ota, Y., Dunkelmann, D., Mittl, P.R.E. and Hilvert, D. (2021) Noncanonical heme ligands steer carbene transfer reactivity in an artificial metalloenzyme. Angew. Chem. Weinheim Bergstr. Ger., 133, 15190–15195.

6. Hayashi, A., Haruna, K.-I., Sato, H., Ito, K., Makino, C., Ito, T. and Sakamoto, K. (2021) Incorporation of Halogenated Amino Acids into Antibody Fragments at Multiple Specific Sites Enhances Antigen Binding. Chembiochem, 22, 120–123.

7. Islam, M., Kehoe, H.P., Lissoos, J.B., Huang, M., Ghadban, C.E., Berumen Sánchez, G., Lane, H.Z. and Van Deventer, J.A. (2021) Chemical Diversification of Simple Synthetic Antibodies. ACS Chem. Biol., 16, 344–359.

8. Costa, S.A., Simon, J.R., Amiram, M., Tang, L., Zauscher, S., Brustad, E.M., Isaacs, F.J. and Chilkoti, A. (2018) Photo-Crosslinkable Unnatural Amino Acids Enable Facile Synthesis of Thermoresponsive Nano-to Microgels of Intrinsically Disordered Polypeptides. Adv. Mater., 30, 1–9.

9. Shapiro, D.M., Mandava, G., Yalcin, S.E., Arranz-Gibert, P., Dahl, P.J., Shipps, C., Gu, Y., Srikanth, V., Salazar-Morales, A.I., O’Brien, J.P., et al. (2022) Protein nanowires with tunable functionality and programmable self-assembly using sequence-controlled synthesis. Nat. Commun., 13, 829.

10. Hershewe, J.M., Wiseman, W.D., Kath, J.E., Buck, C.C., Gupta, M.K., Dennis, P.B., Naik, R.R. and Jewett, M.C. (2020) Characterizing and Controlling Nanoscale Self-Assembly of Suckerin-12. ACS Synth. Biol., 9, 3388–3399.

11. Thyer, R. and Ellington, A.D. (2019) The Role of tRNA in Establishing New Genetic Codes. Biochemistry, 58, 1460–1463.

12. Krahn, N., Tharp, J.M., Crnković, A. and Söll, D. (2020) Engineering aminoacyl-tRNA synthetases for use in synthetic biology. Enzymes, 48, 351–395.

13. Willis, J.C.W. and Chin, J.W. (2018) Mutually orthogonal pyrrolysyl-tRNA synthetase/tRNA pairs. Nat. Chem., 10, 831–837.

14. Dunkelmann, D.L., Willis, J.C.W., Beattie, A.T. and Chin, J.W. (2020) Engineered triply orthogonal pyrrolysyl-tRNA synthetase/tRNA pairs enable the genetic encoding of three distinct non-canonical amino acids. Nat. Chem., 12, 535–544.

15. Cervettini, D., Tang, S., Fried, S.D., Willis, J.C.W., Funke, L.F.H., Colwell, L.J. and Chin, J.W. (2020) Rapid discovery and evolution of orthogonal aminoacyl-tRNA synthetase-tRNA pairs. Nat. Biotechnol., 38, 989–999.

16. Zambaldo, C., Koh, M., Nasertorabi, F., Han, G.W., Chatterjee, A., Stevens, R.C. and Schultz, P.G. (2020) An orthogonal seryl-tRNA synthetase/tRNA pair for noncanonical amino acid mutagenesis in Escherichia coli. Bioorg. Med. Chem., 28, 115662.

17. Italia, J.S., Addy, P.S., Wrobel, C.J.J., Crawford, L.A., Lajoie, M.J., Zheng, Y. and Chatterjee, A. (2017) An orthogonalized platform for genetic code expansion in both bacteria and eukaryotes. Nat. Chem. Biol., 13, 446–450.

18. Dunkelmann, D.L., Oehm, S.B., Beattie, A.T. and Chin, J.W. (2021) A 68-codon genetic code to incorporate four distinct non-canonical amino acids enabled by automated orthogonal mRNA design. Nat. Chem., 13, 1110–1117.

19. Tharp, J.M., Vargas-Rodriguez, O., Schepartz, A. and Söll, D. (2021) Genetic Encoding of Three Distinct Noncanonical Amino Acids Using Reprogrammed Initiator and Nonsense Codons. ACS Chem. Biol., 16, 766–774.

20. Fischer, J.T., Söll, D. and Tharp, J.M. (2022) Directed Evolution of Methanomethylophilus alvus Pyrrolysyl-tRNA Synthetase Generates a Hyperactive and Highly Selective Variant. Frontiers in Molecular Biosciences, 9.

21. Italia, J.S., Addy, P.S., Erickson, S.B., Peeler, J.C., Weerapana, E. and Chatterjee, A. (2019) Mutually Orthogonal Nonsense-Suppression Systems and Conjugation Chemistries for Precise Protein Labeling at up to Three Distinct Sites. J. Am. Chem. Soc., 141, 6204– 6212.

22. Martin, R.W., Des Soye, B.J., Kwon, Y.-C., Kay, J., Davis, R.G., Thomas, P.M., Majewska, N.I., Chen, C.X., Marcum, R.D., Weiss, M.G., et al. (2018) Cell-free protein synthesis from genomically recoded bacteria enables multisite incorporation of noncanonical amino acids. Nat. Commun., 9, 1203.

23. Des Soye, B.J., Gerbasi, V.R., Thomas, P.M., Kelleher, N.L. and Jewett, M.C. (2019) A Highly Productive, One-Pot Cell-Free Protein Synthesis Platform Based on Genomically Recoded Escherichia coli. Cell Chem Biol, 26, 1743-1754.e9.

24. Quast, R.B., Mrusek, D., Hoffmeister, C., Sonnabend, A. and Kubick, S. (2015) Cotranslational incorporation of non-standard amino acids using cell-free protein synthesis. FEBS Lett., 589, 1703–1712.

25. Bundy, B.C. and Swartz, J.R. (2010) Site-specific incorporation of p-propargyloxyphenylalanine in a cell-free environment for direct protein-protein click conjugation. Bioconjug. Chem., 21, 255–263.

26. Worst, E.G., Exner, M.P., De Simone, A., Schenkelberger, M., Noireaux, V., Budisa, N. and Ott, A. (2015) Cell-free expression with the toxic amino acid canavanine. Bioorg. Med. Chem. Lett., 25, 3658–3660.

27. Ozer, E., Chemla, Y., Schlesinger, O., Aviram, H.Y., Riven, I., Haran, G. and Alfonta, L. (2017) In vitro suppression of two different stop codons. Biotechnol. Bioeng., 114, 1065–1073.

28. Cui, Z., Mureev, S., Polinkovsky, M.E., Tnimov, Z., Guo, Z., Durek, T., Jones, A. and Alexandrov, K. (2017) Combining Sense and Nonsense Codon Reassignment for Site-Selective Protein Modification with Unnatural Amino Acids. ACS Synth. Biol., 6, 535–544.

29. Cui, Z., Wu, Y., Mureev, S. and Alexandrov, K. (2018) Oligonucleotide-mediated tRNA sequestration enables one-pot sense codon reassignment in vitro. Nucleic Acids Res., 46, 6387–6400.

30. Lee, K.B., Kim, H.-C., Kim, D.-M., Kang, T.J. and Suga, H. (2012) Comparative evaluation of two cell-free protein synthesis systems derived from Escherichia coli for genetic code reprogramming. J. Biotechnol., 164, 330–335.

31. Lee, K.B., Hou, C.Y., Kim, C.-E., Kim, D.-M., Suga, H. and Kang, T.J. (2016) Genetic Code Expansion by Degeneracy Reprogramming of Arginyl Codons. Chembiochem, 17, 1198–1201.

32. Zimmerman, E.S., Heibeck, T.H., Gill, A., Li, X., Murray, C.J., Madlansacay, M.R., Tran, C., Uter, N.T., Yin, G., Rivers, P.J., et al. (2014) Production of site-specific antibody-drug conjugates using optimized non-natural amino acids in a cell-free expression system. Bioconjug. Chem., 25, 351–361.

33. Borkowski, O., Koch, M., Zettor, A., Pandi, A., Batista, A.C., Soudier, P. and Faulon, J.-L. (2020) Large scale active-learning-guided exploration for in vitro protein production optimization. Nat. Commun., 11, 1872.

34. Zawada, J.F., Yin, G., Steiner, A.R., Yang, J., Naresh, A., Roy, S.M., Gold, D.S., Heinsohn, H.G. and Murray, C.J. (2011) Microscale to manufacturing scale-up of cell-free cytokine production--a new approach for shortening protein production development timelines. Biotechnol. Bioeng., 108, 1570–1578.

35. Garenne, D., Thompson, S., Brisson, A., Khakimzhan, A. and Noireaux, V. (2021) The all-E. coliTXTL toolbox 3.0: new capabilities of a cell-free synthetic biology platform. Synth. Biol., 6, ysab017.

36. Albayrak, C. and Swartz, J.R. (2013) Cell-free co-production of an orthogonal transfer RNA activates efficient site-specific non-natural amino acid incorporation. Nucleic Acids Res., 41, 5949–5963.

37. Wan, W., Huang, Y., Wang, Z., Russell, W.K., Pai, P.-J., Russell, D.H. and Liu, W.R. (2010) A facile system for genetic incorporation of two different noncanonical amino acids into one protein in Escherichia coli. Angew. Chem. Int. Ed Engl., 49, 3211–3214.

38. Zubi, Y.S., Seki, K., Li, Y., Hunt, A.C., Liu, B., Roux, B., Jewett, M.C. and Lewis, J.C. (2022) Metal-responsive regulation of enzyme catalysis using genetically encoded chemical switches. Nat. Commun., 13, 1–10.

39. Swartz, J.R., Jewett, M.C. and Woodrow, K.A. (2004) Cell-Free Protein Synthesis With Prokaryotic Combined Transcription-Translation. In Balbás, P., Lorence, A. (eds), Recombinant Gene Expression: Reviews and Protocols. Humana Press, Totowa, NJ, pp. 169–182.

40. Kightlinger, W., Duncker, K.E., Ramesh, A., Thames, A.H., Natarajan, A., Stark, J.C., Yang, A., Lin, L., Mrksich, M., DeLisa, M.P., et al. (2019) A cell-free biosynthesis platform for modular construction of protein glycosylation pathways. Nat. Commun., 10, 5404.

41. Sun, Z.Z., Yeung, E., Hayes, C.A., Noireaux, V. and Murray, R.M. (2014) Linear DNA for rapid prototyping of synthetic biological circuits in an Escherichia coli based TX-TL cell-free system. ACS Synth. Biol., 3, 387–397.

42. Jewett, M.C., Calhoun, K.A., Voloshin, A., Wuu, J.J. and Swartz, J.R. (2008) An integrated cell-free metabolic platform for protein production and synthetic biology. Mol. Syst. Biol., 4, 220.

43. Fechter, P., Rudinger, J., Giegé, R. and Théobald-Dietrich, A. (1998) Ribozyme processed tRNA transcripts with unfriendly internal promoter for T7 RNA polymerase: production and activity. FEBS Lett., 436, 99–103.

44. Xie, J., Liu, W. and Schultz, P.G. (2007) A genetically encoded bidentate, metal-binding amino acid. Angew. Chem. Int. Ed Engl., 46, 9239–9242.

45. Bryson, D.I., Fan, C., Guo, L.-T., Miller, C., Söll, D. and Liu, D.R. (2017) Continuous directed evolution of aminoacyl-tRNA synthetases. Nat. Chem. Biol., 13, 1253–1260.

46. Chatterjee, A., Sun, S.B., Furman, J.L., Xiao, H. and Schultz, P.G. (2013) A versatile platform for single- and multiple-unnatural amino acid mutagenesis in Escherichia coli. Biochemistry, 52, 1828–1837.

47. Ryu, Y. and Schultz, P.G. (2006) Efficient incorporation of unnatural amino acids into proteins in Escherichia coli. Nat. Methods, 3, 263–265.

48. Aerni, H.R., Shifman, M.A., Rogulina, S., O’Donoghue, P. and Rinehart, J. (2015) Revealing the amino acid composition of proteins within an expanded genetic code. Nucleic Acids Res., 43, e8.

49. Wan, W., Tharp, J.M. and Liu, W.R. (2014) Pyrrolysyl-tRNA synthetase: an ordinary enzyme but an outstanding genetic code expansion tool. Biochim. Biophys. Acta, 1844, 1059– 1070.

50. Nozawa, K., O’Donoghue, P., Gundllapalli, S., Araiso, Y., Ishitani, R., Umehara, T., Söll, D. and Nureki, O. (2009) Pyrrolysyl-tRNA synthetase-tRNA(Pyl) structure reveals the molecular basis of orthogonality. Nature, 457, 1163–1167.

51. Bednar, R.M., Jana, S., Kuppa, S., Franklin, R., Beckman, J., Antony, E., Cooley, R.B. and Mehl, R.A. (2021) Genetic Incorporation of Two Mutually Orthogonal Bioorthogonal Amino Acids That Enable Efficient Protein Dual-Labeling in Cells. ACS Chem. Biol., 16, 2612–2622.

52. Ostrov, N., Landon, M., Guell, M., Kuznetsov, G., Teramoto, J., Cervantes, N., Zhou, M., Singh, K., Napolitano, M.G., Moosburner, M., et al. (2016) Design, synthesis, and testing toward a 57-codon genome. Science, 353, 819–822.

53. Fredens, J., Wang, K., de la Torre, D., Funke, L.F.H., Robertson, W.E., Christova, Y., Chia, T., Schmied, W.H., Dunkelmann, D.L., Beránek, V., et al. (2019) Total synthesis of Escherichia coli with a recoded genome. Nature, 569, 514–518.

